# A novel gyrovirus associated with a fatal respiratory disease in yellow-eyed penguin (*Megadyptes antipodes*) chicks

**DOI:** 10.1101/2022.09.15.508173

**Authors:** Janelle R. Wierenga, Kerri J. Morgan, Stuart Hunter, Harry S. Taylor, Lisa S. Argilla, Trudi Webster, Jeremy Dubrulle, Fátima Jorge, Mihnea Bostina, Laura Burga, Edward C. Holmes, Kate McInnes, Jemma L. Geoghegan

**Author notes:** Corresponding author: Jemma L. Geoghegan.

## Abstract

Yellow-eyed penguins (*Megadyptes antipodes*), or *hoiho* in te reo Māori, are predicted to become extinct on mainland Aotearoa New Zealand in the next few decades, with infectious disease a significant contributor to their decline. A recent disease phenomenon termed respiratory distress syndrome (RDS) causing lung pathology has been identified in very young chicks. To date, no causative pathogens for RDS have been identified. In 2020 and 2021, the number of chick deaths from suspected RDS increased four- and five-fold, respectively, with a mortality rate of >90%. Here, we aimed to identify possible pathogens responsible for RDS disease impacting yellow-eyed penguins. Total RNA was extracted from tissue samples collected during post-mortem of 43 chicks and subject to metatranscriptomic sequencing. From these data we identified a novel and highly abundant gyrovirus in 80% of tissue samples. This virus exhibited only 41% amino acid identity within VP1 to its closest relative, *Gyrovirus 8*, discovered in a diseased seabird. No other exogenous viral transcripts, nor pathogenic bacterial, protozoal and fungal organisms, were identified in these tissues. Due to the high relative abundance of viral reads, it is likely that this novel gyrovirus is associated with RDS in yellow-eyed penguin chicks.

**Author Summary:** New Zealand’s population of yellow-eyed penguins, also called *hoiho*, are predicted to become extinct in the next 20-30 years, with disease a major factor contributing to their decline. A new disease, causing fluid and bleeding into the lungs, was initially identified in 2019 in very young chicks. It was characterised as causing respiratory distress with a mortality of >90% usually within the first week of life. To date, no causative pathogens of the disease have been identified. We aimed to identify possible pathogens responsible for respiratory disease in these penguin chicks. A metatranscriptomic survey of dead chicks identified a novel and highly abundant gyrovirus present in diseased tissue, with closely related viruses causing disease in other avian hosts. It is, therefore, highly likely that this novel gyrovirus is associated with respiratory disease in these chicks. This finding offers the potential to increase the success of disease management in the critically endangered yellow-eyed penguin and possibly other at-risk penguin species. The potential to lessen mortality and slow the decline of the species is essential in protecting the biodiversity of New Zealand’s fauna and flora.

## Introduction

Yellow-eyed penguins (*Megadyptes antipodes*), or *hoiho* in te reo Māori, are a critically endangered species endemic to Aotearoa New Zealand and considered one of the rarest penguins in the world. There are two populations of yellow-eyed penguins: northern (occupying mainland New Zealand and Stewart Island/Rakiura plus adjacent islands) and southern (on subantarctic Auckland Islands and Campbell Island). The northern population has experienced a dramatic decline with numbers decreasing by 75% over the past 30 years. Due to this decline, it is speculated that yellow-eyed penguins will become extinct from the mainland of New Zealand within the next two decades (1).

Infectious diseases affecting young chicks are one of the major threats to the extinction of yellow-eyed penguins. Diphtheritic stomatitis associated with *Corynebacterium* spp. has caused significant mortality since its emergence in 2002, with one study reporting the disease in 50% of chicks examined post-mortem (2). Although its role in chick mortality is unclear, *Leucocytozoon* spp. have been reported at a high prevalence and causing mortality in both mainland and subantarctic populations (3). Another disease, termed respiratory distress syndrome (RDS), was initially identified in 2019, although historical epidemiological records show the first suspected cases as early as 2015. This manifests as lung congestion and haemorrhage along with lymphoid depletion in the spleen and bursa (unpublished). In 2020 and 2021, the number of chick deaths from RDS increased four- and five-fold compared to 2019, respectively, with a mortality rate of over 90%, and chicks typically succumbed to the disease within the first week of life.

We used a metatranscriptomic approach to identify possible causative agents associated with RDS in yellow-eyed penguins. This led to the discovery of a novel and highly abundant gyrovirus. Gyroviruses are small, non-enveloped DNA viruses within the family *Anelloviridae*, initially characterised as circoviruses but reclassified into the family in 2017 (4, 5). Currently classified gyroviruses possess negative-sense, single-stranded DNA circular genomes of approximately 2000-2400 nucleotides in length. Gyroviruses utilise the host machinery for replication and have three overlapping viral protein (VP) open-reading frames: VP1, that encodes the viral capsid; VP2; and VP3.

Gyroviruses are known to cause disease in avian hosts, while additional viral species have been identified from the faeces of domestic cats (6), mice (7), ferrets (8), dogs (9) and humans (10). The best-known virus in this genus is *Chicken anaemia virus* (CAV), which causes anaemia, poor growth and severe immunosuppression in young chicks (11–14). CAV is found worldwide and causes significant mortality among chicks not protected by maternal antibodies. Other gyroviruses have been identified in diseased avian hosts, including in seabirds (15–19). Herein, we describe a new gyrovirus associated with RDS in yellow-eyed penguins.

## Results

### Disease Investigation, Histology and Electron Microscopy

Yellow-eyed penguin chicks that died with suspected RDS during the November to December 2021 hatching season across eastern Otago, New Zealand were subject to investigation to identify the possible causative agent of the disease. These chicks typically died within the first week of life with ages ranging between two and 10 days old (median = five days). A total of 43 penguin chicks underwent post-mortem examination within 1-2 days of death after showing possible signs of RDS.

From a total of 137 wild yellow-eyed penguin chicks that were admitted to the Dunedin Wildlife Hospital during the 2021 breeding season, 31 (22.6%) that presented between 3-19 November demonstrated clinical evidence of respiratory disease within their first week of life. The majority of these birds died (n=24, 77%) or were euthanised (n=3, 10%) within 12-24 hours of presentation, with one bird (3%) surviving for 5 days with intensive veterinary support before succumbing. Three (10%) birds were successfully treated and were returned to the nest.

In most cases, the clinical progression of respiratory disease was very rapid. Generally, affected birds demonstrated a progressive increase in respiratory rate and effort, and as the disease progressed, birds became very weak and often recumbent, with pale mucous membranes and evidence of hypothermia despite the provision of external heat. Terminally, chicks presented with coelomic distension, presumably due to overinflation of airsacs as a result of agonal gasping, and birds were visibly cyanotic with a reduced level of consciousness. In addition to the 28 birds that died in hospital, one died enroute to hospital, one was euthanised due to a limb deformity and a further 13 neonates died at the nest during this same period (total n=43).

Gross post-mortem examination identified abnormal lung tissue bilaterally in 88% (38/43) of chicks, characterised as dark pink to dark purple lungs with obvious haemorrhage and clots within the lungs. A consistent histopathological finding was the presence of proteinaceous fluid (Figure 1a, red star) and haemorrhage (Figure 1a, yellow stars) within the parabronchi combined with partial or full collapse of the smaller airways, the atria and air-capillaries (Figure 1). Epithelial cells lining the atria and air-capillaries were often hyperplastic and hypertrophic and there was an increased presence of macrophages within these airways. Generalised congestion as well as oedema of interlobular septae was also a consistent feature. Within the spleen and bursa, there was lymphoid depletion as well as evidence of active lympholysis.

**Figure 1.**
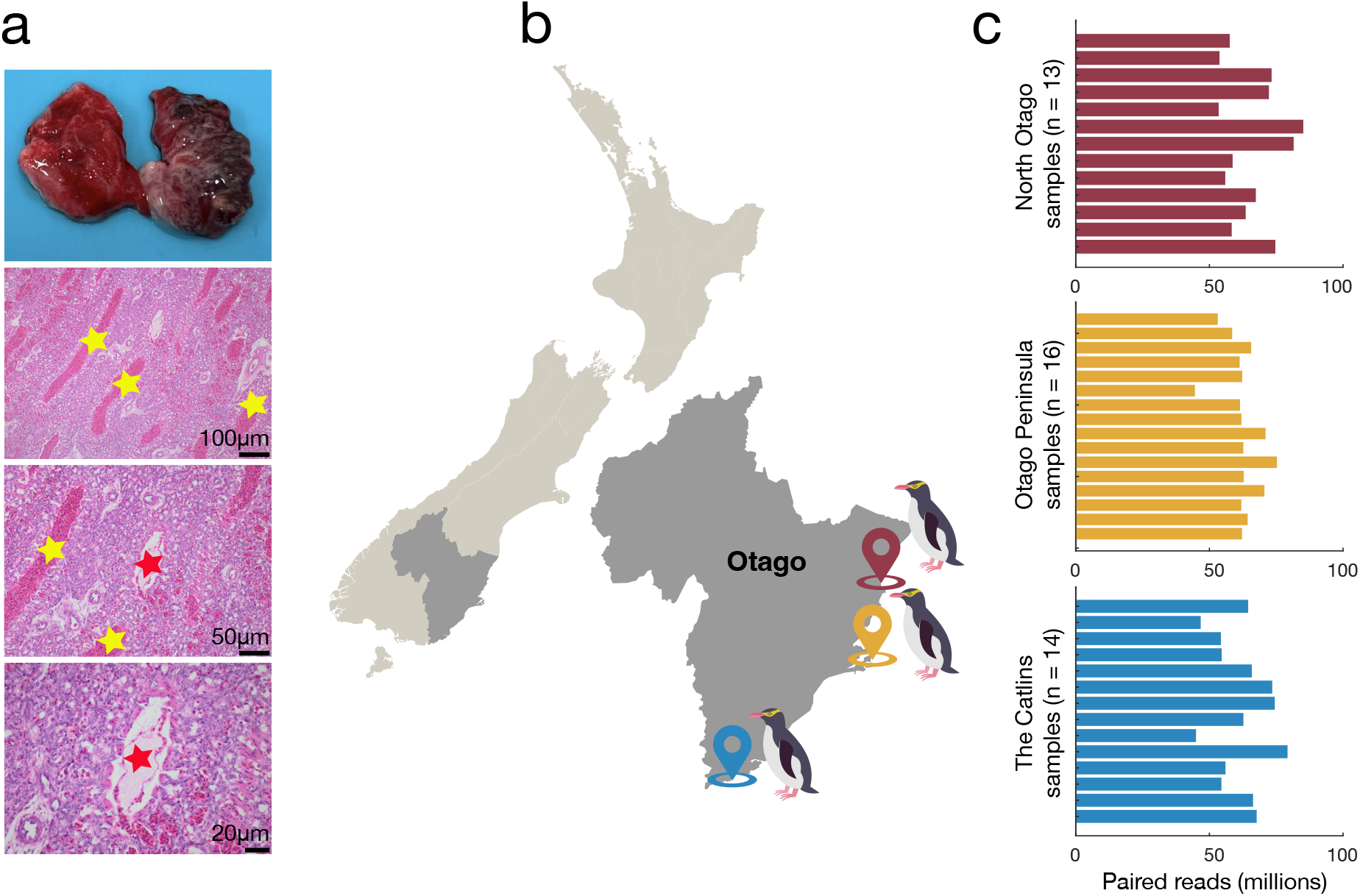
(a) Fresh lung tissue (top) and lung histology from a yellow-eyed penguin (*Megadyptes antipodes*) chick that was diagnosed with respiratory distress syndrome (RDS); yellow stars: blood within parabronchi, red stars: proteinaceous fluid within parabronchi. (b) Sampling locations of yellow-eyed penguin chicks across the east coast of Otago, New Zealand (north Otago = red; Otago peninsula = yellow; The Catlins = blue). (c) Total number of paired-end RNA sequencing reads obtained from yellow-eyed penguin chick tissue samples with ribosomal RNA depletion.

Yellow-eyed penguin chicks (n=9) were characterised as ‘RDS suspected’ when they had clinical signs associated with respiratory disease prior to death and/or upon gross post-mortem the lungs were characterised as dark pink to purple in colour with or without clots in the lung tissue upon extraction. Chicks (n=31) were characterised as ‘RDS confirmed’ based on histopathological analyses as described above. Similarly, chicks (n=3) were characterised as ‘RDS not suspected’ based upon histopathological analyses of the tissues. Of the three chicks without RDS, the cause of death for two were consistent with aspiration pneumonia at 10 and 11 days of age; while the cause of death for the remaining chick (15 days of age at time of death) was unknown.

Electron microscopy investigation of the diseased lung and spleen tissue demonstrated extended cytopathic effect in all the specimens (Supplementary Figures 1 and 2). This included large vacuoles, swollen mitochondria, and nuclear fragmentation with chromatin aggregates close to the nuclear rim. Notably, cells in the spleen sample showed characteristic electron dense rings in the nuclei similar to reports of CAV (20).

### Viral Diversity and Abundance

Metatranscriptomic sequencing of multiple tissue samples (bursa, kidney, liver, lung, spleen) from each individual yielded sequencing libraries containing between 44.5 - 85.1 million paired-end reads that were *de novo* assembled into 231,069 - 964,934 contigs (Figure 1). Forty-three libraries were evaluated for viral transcripts likely infecting eukaryotic hosts (excluding plant, fungi and protists). A sequence similarity-based search for exogenous viruses identified viral transcripts as a novel gyrovirus (*Anelloviridae*) in 34 of the 43 samples (79%), tentatively named yellow-eyed penguin gyrovirus. No other exogenous viral transcripts were identified. Of those samples with suspected and confirmed RDS, gyrovirus was identified in 32 of 40 (80%) at standardised abundances ranging from 7 x 10^-7^ to 0.00015, comparable to other avian viral abundances found in penguins (21) (Figure 2, Supplementary Table 1). Two out of three samples that were not suspected of having RDS upon gross post-mortem also contained gyrovirus sequencing reads at similar abundances, although this could reflect subjective aspects of disease diagnosis (see Discussion).

**Figure 2.**
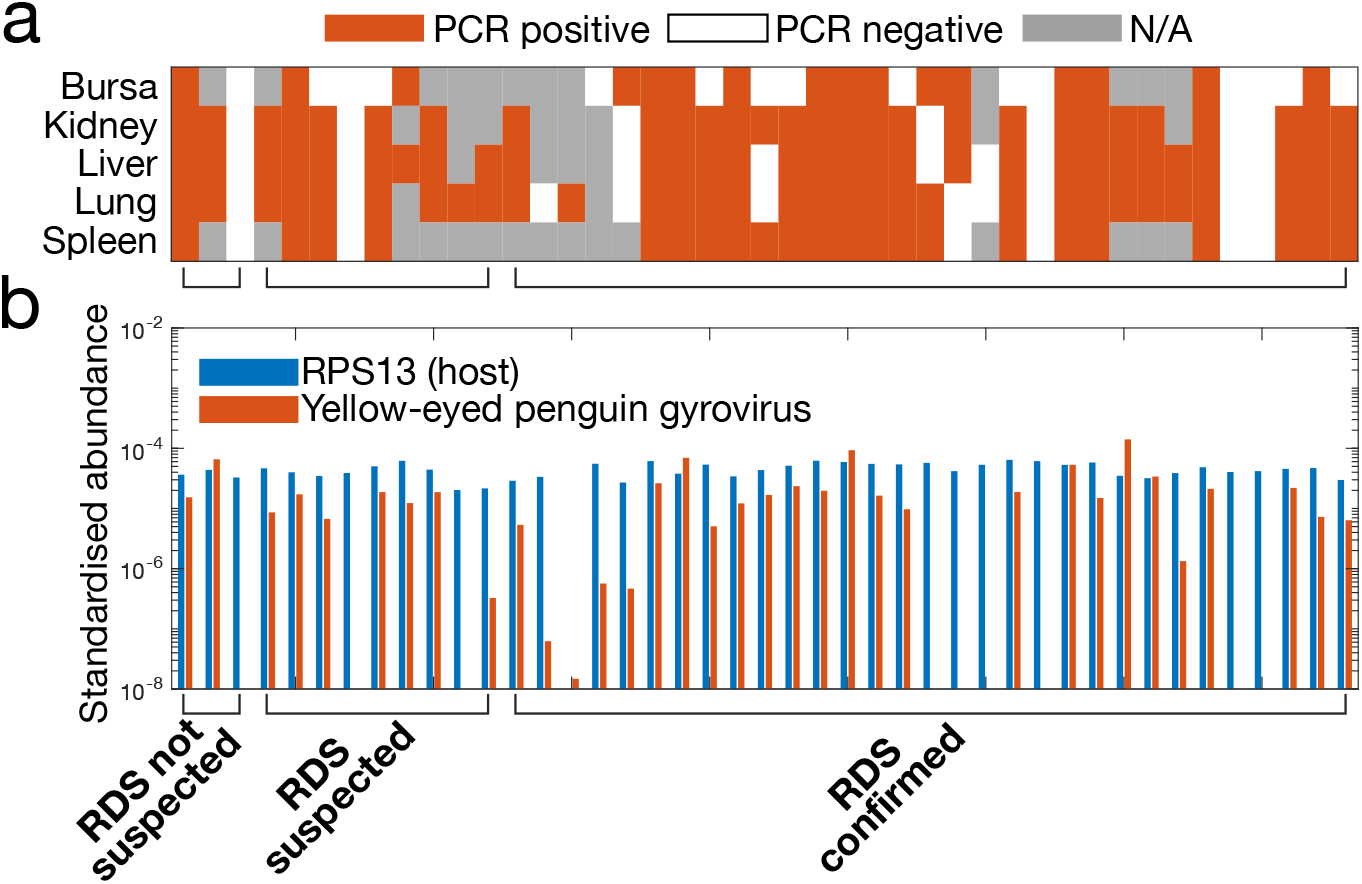
(a) Tissue samples collected from deceased yellow-eyed penguin chicks (*Megadyptes antipodes*) during the 2021 breeding season that were PCR positive for a novel yellow-eyed penguin gyrovirus (red), PCR negative (white) or not collected (or exhausted) (grey) across five different tissue samples. (b) Standardised relative abundances of the pooled RNA sequencing libraries of a host gene, ribosomal protein s13 (RPS13) (blue) and yellow-eyed penguin gyrovirus (red) across samples that were deemed: (i) respiratory disease syndrome (RDS) not suspected; (ii) RDS suspected at gross post-mortem; and (iii) RDS confirmed with histology.

### PCR Confirmation of Yellow-Eyed Penguin Gyrovirus

PCR was performed to confirm and identify the presence of yellow-eyed penguin gyrovirus from the same tissue samples as used for the metatranscriptomic sequencing (Figure 2). The PCR was confirmed and optimised on two samples that had confirmed RDS status upon histopathological examination and high relative abundances of yellow-eyed penguin gyrovirus sequencing reads.

PCR was then performed on all samples as individual tissues (including bursa, kidney, liver, lung and spleen). Three samples that tested positive upon PCR for yellow-eyed penguin gyrovirus and were suspected or confirmed as RDS had no gyrovirus sequencing reads (Figure 2). PCR results largely agreed with the presence or absence of yellow-eyed penguin gyrovirus in the pooled RNA sequencing results. Where discrepancies occurred, for example those samples that were PCR positive, but no virus reads were detected, were likely due to the pooled RNA obtained from all tissue samples diluting the overall RNA in the sample coupled with high sensitivity of PCR.

### Genome Organisation and Phylogenetic Characterisation of a Novel Gyrovirus

A total of 29 full and a further five partial genomes of the novel virus were assembled *de novo* from the 43 samples, which all shared >95% nucleotide sequence identity. The novel yellow-eyed penguin gyrovirus genome contained 2610 nucleotides, the largest known gyrovirus genome, and three overlapping open reading frames: VP1, VP2 and VP3 (Figure 3). Both VP1 and VP2 shared 41% and 27% amino acid identity, respectively, to its closest relative, *Gyrovirus 8*, identified in a diseased northern fulmar (*Fulmarus glacialis*) sampled from California (15). In contrast, VP3 did not share sequence homology to any other known protein, as is often the case with this ORF in other gyroviruses. The VP1 of the yellow-eyed gyrovirus contained the signature ‘WW[R/N]W[S/A]’ motif at position 153 with an arginine-rich N-terminus region, while the VP2 contained the conserved ‘WX7HX3CXCX5H’ motif at residue 96 (Figure 3).

**Figure 3.**
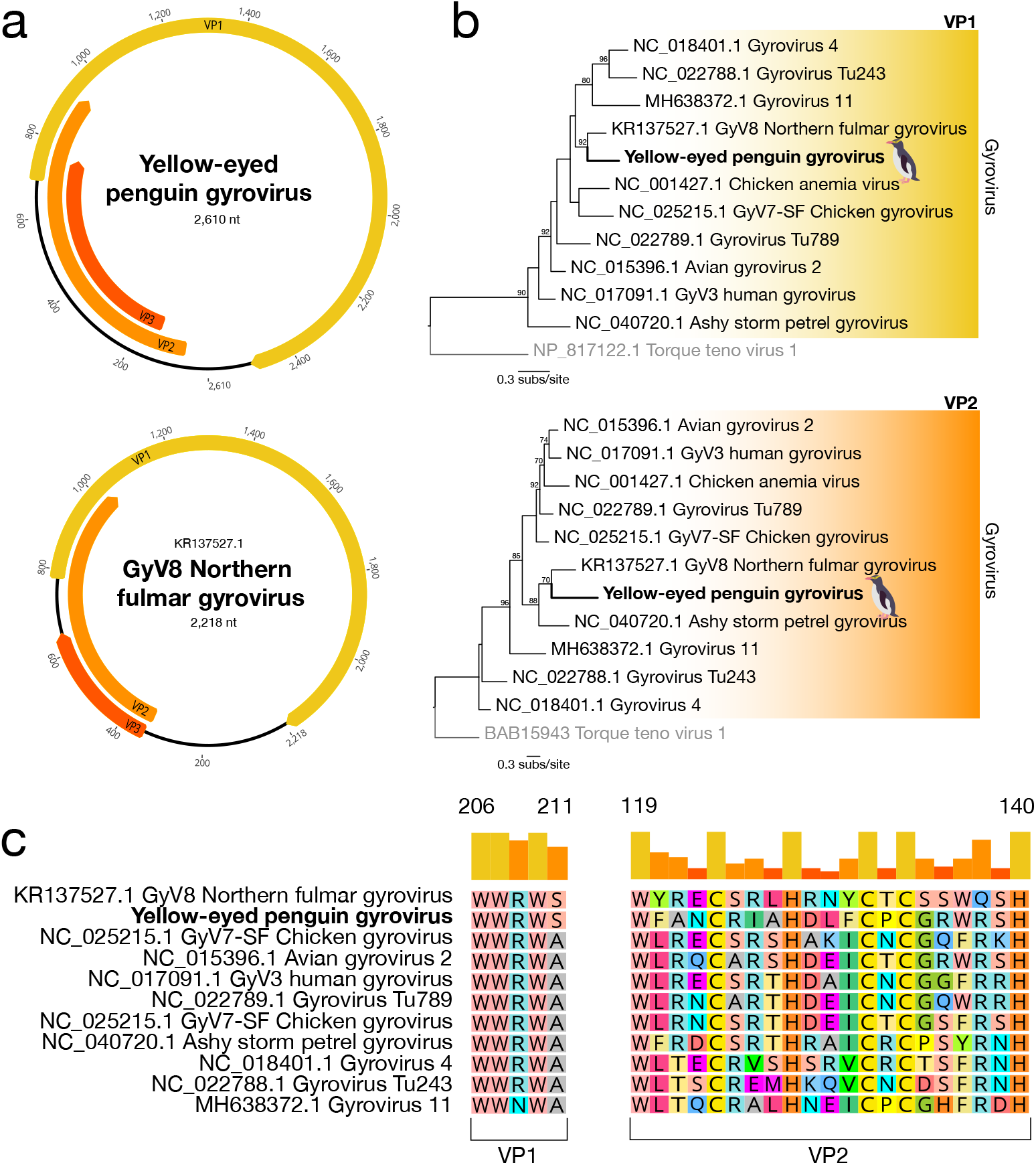
(a) Genome structure of yellow-eyed penguin gyrovirus (top) illustrating a monopartite, circular, ssDNA genome containing 2610 nucleotides, in comparison with Gyrovirus 8 identified in a northern fulmar (*Fulmarus glacialis*) from California (lower). The positions of viral proteins (VP) 1, 2 and 3 are shown. (b) Maximum likelihood phylogenetic trees of VP1 (top) and VP2 (lower) of viruses within the Gyrovirus genus using Torque teno virus 1 as an outgroup. The topological position of yellow-eyed penguin gyrovirus is shown in bold. Branches are scaled to the number of amino acid substitutions per site. Node support of >70% is shown. (c) Signature conserved motifs within VP1 and VP2 across members of the Gyrovirus genus.

To infer the evolutionary history of yellow-eyed penguin gyrovirus, we estimated phylogenetic trees for both VP1 and VP2 among other known gyroviruses using *Torque teno virus 1*, a nongyrovirus member of the *Anelloviridae*, as an outgroup (Figure 3). Within VP1, yellow-eyed gyrovirus clustered with *Gyrovirus 8*, which caused disease in the northern fulmar, while VP2 fell within a clade of gyroviruses infecting seabirds, including *Ashy storm-petrel gyrovirus* (16).

We next inferred the phylogeography of all full and partial yellow-eyed gyrovirus genomes obtained (34 of 43 samples) across the three sampling sites (Figure 4). All genomes fell across two distinct genomic clades (termed A and B), that showed little spatial structure with all three sampling locations contained in both clades. We also found no clustering by gross or confirmed post-mortem RDS status.

**Figure 4.**
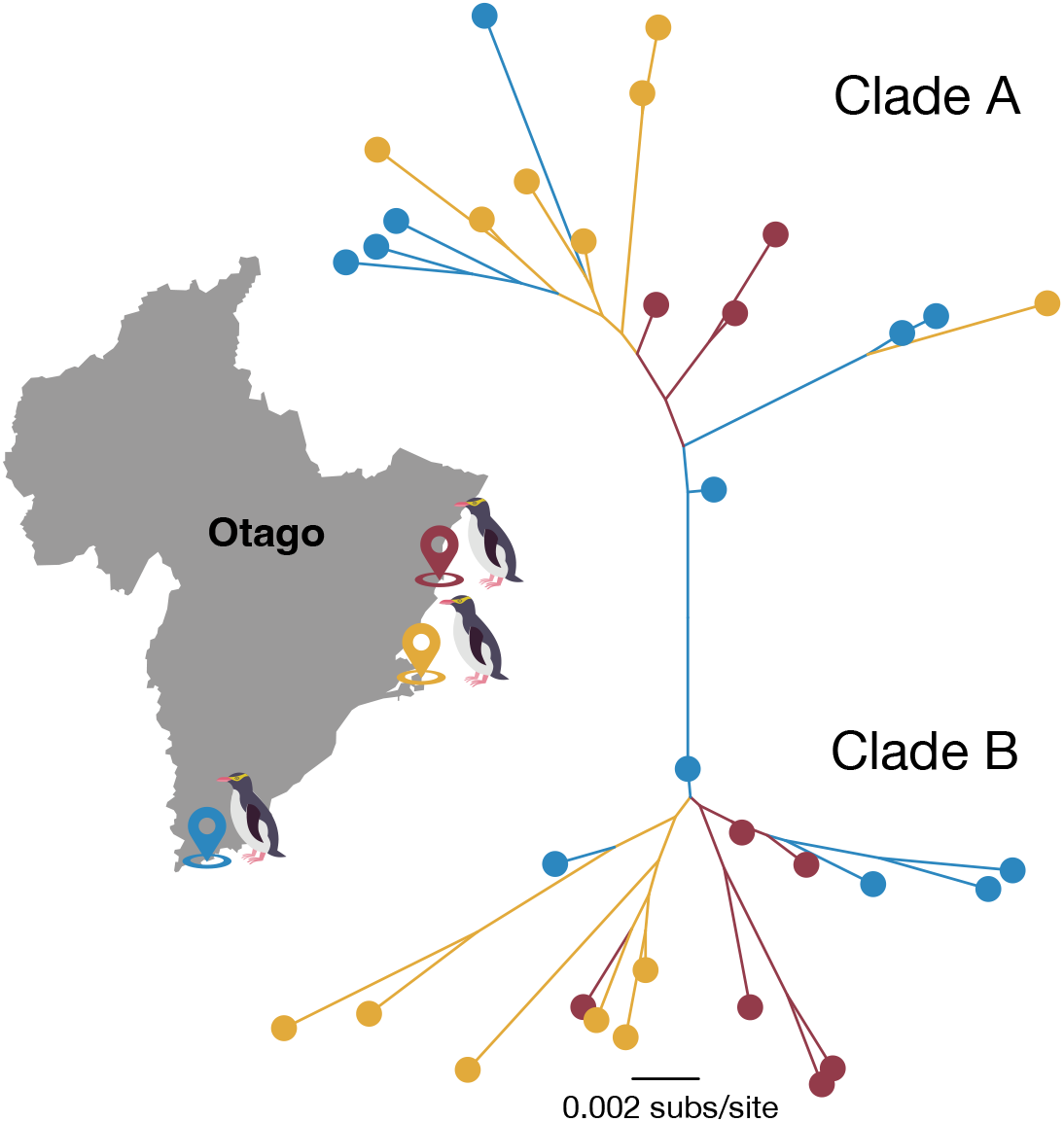
Unrooted maximum likelihood phylogenetic tree of the full (n = 29) or partial (n = 5) genomes of yellow-eyed penguin gyrovirus identified from multiple breeding sites on mainland New Zealand in organ tissue metatranscriptomes. Branches are scaled to the number of nucleotide substitutions per site. Branches and tips are coloured to illustrate the sampling site shown on the adjacent map. All samples fell within two major clades, annotated A and B.

### Total Infectome Characterisation

RNA sequencing data were further evaluated for the presence of bacterial, protozoal and fungal organisms (Figure 5). The *Enterobacteriaceae*, *Enterococcaceae* and *Staphylococcaceae* families were consistently the most abundant and prevalent across samples while *Aspergillus* spp. were the most abundant fungi detected. Transcripts identified as *Plasmodiaaceae* were found in very few of the libraries in low abundances. It is worth noting, however, that taxonomic misassignment of host genes to microbial organisms is commonplace. In marked contrast to yellow-eyed penguin gyrovirus, no bacterial, protozoal or fungal organism had a clear association with RDS.

**Figure 5.**
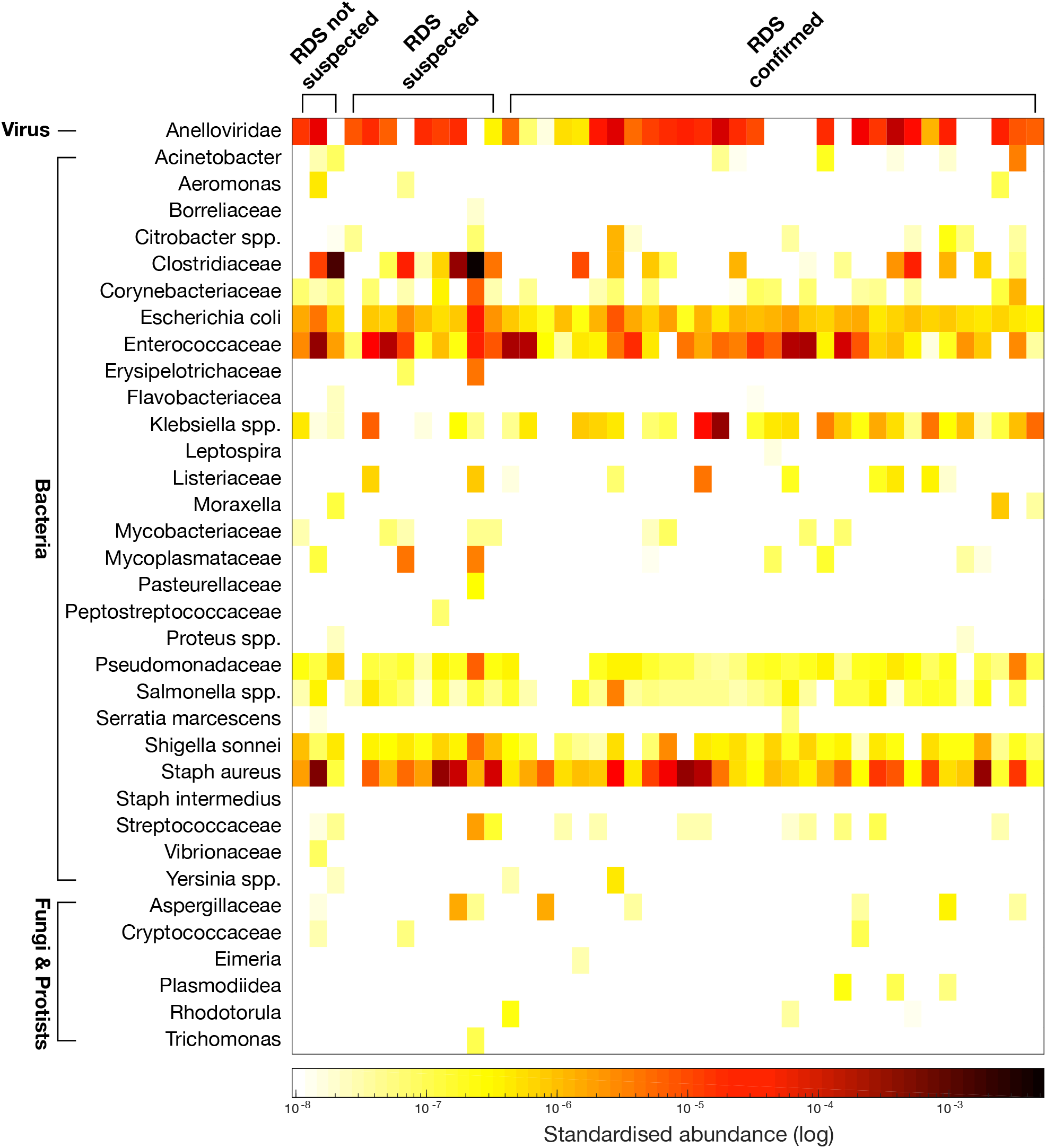
Standardised read abundance of viral, bacteria, fungi and protist families/genera detected through metatranscriptomic sequencing of tissues from deceased yellow-eyed penguin (*Megadyptes antipodes*) chicks (n=43) across samples that were deemed: (i) respiratory disease syndrome (RDS) not suspected; (ii) RDS suspected at gross post-mortem; and (iii) RDS confirmed with histology.

## Discussion

Our metatranscriptomic investigation into the possible causative agents of the highly fatal respiratory disease syndrome (RDS) affecting yellow-eyed penguin chicks revealed a novel, divergent and highly abundant gyrovirus within the *Anelloviridae*, tentatively named yellow-eyed penguin gyrovirus. This novel gyrovirus was most closely related to *Gyrovirus 8* found in a diseased northern fulmar (15). Other viruses within the genus *Gyrovirus* are known to infect avian species and cause similar pathology. In particular, *Chicken anaemia virus* (CAV) is a well-characterised pathogenic virus in the *Anelloviridae* causing anaemia, haemorrhage and immunosuppression in poultry chicks, inflicting significant economic losses to the industry (13, 22–24). CAV can be transmitted both horizontally and vertically and, as a small circular DNA virus, is highly stable and resistant to most disinfectants (13, 25–27).

Phylogenetic analysis of full and partial yellow-eyed penguin gyrovirus genomes identified two distinct clades with little spatial structure among sampling locations and no clustering with gross post-mortem RDS status. The relatively high diversity among yellow-eyed penguin gyrovirus genomes suggested sustained transmission of the virus in the population, compatible with outbreaks dating to at least 2015 when the disease was first identified during post-mortem examination. Further testing and genome sequencing of historical samples suspected of RDS will help to elucidate the evolutionary history of this novel virus.

RDS in yellow-eyed penguin chicks is a recently identified disease causing high mortality despite intensive veterinary intervention. During the 2020 and 2021 seasons, 15% and 23% of the chicks that were monitored on the mainland died as a result of RDS, respectively. Although clinical symptoms are highly suggestive, there are a number of potential causes of respiratory disease in young penguins (2, 28–31). At present, RDS is highly suspected on post-mortem with grossly visible dark pink to purple lungs with extensive haemorrhage identifiable on cut surface and histological examination provides further confirmation. As a consequence, we developed a PCR test for yellow-eyed penguin gyrovirus that can be used in future breeding seasons to understand the true prevalence of the virus among both diseased and healthy populations.

We attempted to identify other potential causative pathogens of bacterial, protozoal or fungal origin. The majority of bacterial organisms identified were likely typical commensal flora rather than pathogens. In addition, it is likely that much of the bacterial growth occurred post-mortem: dead chicks were in warm environments either in hospital incubators or under the parent in the nest, which could have promoted growth of bacteria in tissue that is normally sterile. Further, contamination during gross post-mortem from the gastrointestinal tract was also possible.

A few samples that tested positive upon PCR for yellow-eyed gyrovirus contained no gyrovirus sequencing reads. In addition, several samples that were confirmed or suspected RDS had an absence of gyrovirus reads. Nevertheless, the range of possible disease phenotypes for RDS is not well understood and gross post-mortem is highly subjective. In addition, there were inconsistencies of tissue types collected across individuals as well as possible tissue decomposition that would likely affect nucleic acid extraction. Finally, pooled RNA from across the five tissue types collected in most cases likely diluted viral RNA that was present in only small abundances and/or in few tissues.

Overall, both the prevalence and relatively high abundance of yellow-eyed gyrovirus supports the pathogenic potential of this virus. In addition, the absence of other exogenous viral transcripts and other known pathogenic microbes suggests that yellow-eyed penguin gyrovirus is likely associated with RDS. Further work is necessary to confirm this association, to fully exclude yellow-eyed gyrovirus as an opportunistic secondary infection and to better understand the characteristics of this disease (e.g. transmission) to enable appropriate management recommendations.

## Materials & Methods

### Ethics

This study was conducted under animal ethics approval MUAEC Protocol 21/42 from Massey University and the New Zealand Department of Conservation Permit Authorisation Number 94843-FAU and 94902-FAU.

### Sample Collection and Post-Mortem

Three populations of yellow-eyed penguins are monitored annually by the New Zealand Department of Conservation along with various organisations from three regions across Otago, New Zealand. Samples collected in this study originated from Moeraki, North Otago (GPS −45° 21’59.99” S,170° 50’ 59.99” E; n = 13), Otago Peninsula (−45° 51’ 17.99” S, 170° 38’ 59.99” E; n = 16) and The Catlins (−46° 29’ 59.99” S, 169° 29’ 59.99” E; n = 14) during the November and December 2021 breeding season (Figure 1). Newly hatched chicks were monitored and weighed every two to three days and chicks from nests that demonstrated respiratory symptoms or weight loss were immediately transferred to the Dunedin Wildlife Hospital for veterinary treatment.

Affected chicks were immediately placed in incubators set at 32°C (34-35°C for those with hypothermia) flooded with 100% oxygen. Lactated Ringer’s solution was administered subcutaneously (100ml/kg/day) and broad spectrum anti-microbials were initiated including 100mg/kg amoxyclav (400mg amoxycillin and clavulanic acid 57mg) PO BID; 15mg/kg enrofloxacin PO BID and 5mg/kg voriconazole PO SID-BID. Meloxicam (5mg/kg PO) was provided once birds were rehydrated. As chicks generally presented with ileus due to being hypothermic and clinically unwell, food was withheld until chicks clinically stabilised, after which fish slurry was provided via gavage up to five times daily as tolerated.

Neonatal yellow-eyed penguin chicks that died during this period had gross post-mortem examinations performed by a veterinarian within 24 hours of death, with dead chicks placed in a refrigerator at 4°C prior to examination. Tissues that were collected included brain, entire gastrointestinal tract including the oral cavity, lung, heart, thyroid glands, liver, spleen, bursa of Fabricius, adrenal glands, kidneys, yolk sac (if present) and skeletal muscle, although not all tissues were collected for all individuals (see Supplementary Table 1). In situations of marked decomposition of the body either the lung tissue, as this was the observable abnormal tissue that could still be evaluated grossly, or the lung, liver and kidney tissue were collected for analysis; however, in most cases, lung, liver, kidney, bursa of Fabricus and spleen were utilised. Tissues were stored in sterile vials in RNAlater^®^ as well as in 10% formalin for histopathological analyses. Tissues in RNAlater^®^ were immediately stored in a −80°C freezer.

### Histology

Formalin fixed tissue samples from 34 chicks (out of 43 total) were embedded in paraffin and 4-μm thick sections were stained with hematoxylin and eosin for histological examination. Tissues were examined for the presence of proteinaceous fluid and haemorrhage within the parabronchi combined with partial or full collapse of the smaller airways, the atria and air-capillaries in the lungs.

### Electron Microscopy

Initially small lung tissue fragments collected during post mortem examination were fixed in freshly prepared 2% glutaraldehyde in 0.1M cacodylate buffer pH7.4 for at least 24 hours. Following identification of yellow-eyed penguin gyrovirus by metatranscriptomic sequencing in most tissue samples, spleen samples also collected during necropsy and stored at −80°C were thawed and fixed as described above. The following steps were performed at room temperature under agitation. After primary fixation, tissues were washed three times for 10 minutes with 0.1M cacodylate buffer and post-fixed in 2% osmium tetroxide in 0.1M cacodylate buffer for one hour. Specimens were then washed three times for 10 minutes with 0.1M cacodylate buffer, then twice with doubled distilled water for 10 minutes followed by dehydration through a series of ascending grades of ethanol concentrations (50%, 70% and 95%, and 2 x 100% for 10 minutes each, and finally 100% for 20 minutes), and rinsed twice with propylene oxide for 15 minutes. The propylene oxide was replaced by dilution of Suprr’s resin in propylene oxide (1:1 propylene oxide:resin for 20 minutes and 1:2 propylene oxide:resin for 40 minutes). Tissue samples transitioned through three Spurr’s resin changes with four-hour incubation time. Each fragment was then transferred to an embedding mould with fresh resin and polymerised for 48 hours at 60°C. Ultrathin sectioning (85nm) was done with a diamond knife on a Leica EM UC7 ultramicrotome and collected on formvar-coated copper slot grids. Sections were stained with uranyl acetate and lead citrate and investigated using a Philips CM100 transmission electron microscope (TEM).

### RNA Extraction and Sequencing

Frozen tissue was partially thawed and submerged in lysis buffer containing 1% β-mercaptoethanol and 0.5% Reagent DX (Qiagen) before tissues were homogenised together with TissueRupture (Qiagen). The homogenate was centrifuged to remove any potential tissue residues, and RNA from the clear supernatant was extracted using the Qiagen RNeasy Plus Mini Kit. RNA was quantified using NanoDrop (ThermoFisher) and RNA from each tissue type was pooled into one sample per individual. Extracted RNA was subject to total RNA sequencing. Libraries were prepared using the Illumina Stranded Total RNA Prep with Ribo-Zero Plus (Illumina). Paired-end 150bp sequencing of the RNA libraries was performed on the Illumina NovaSeq 6000 platform.

### Virome Composition Analysis

Sequencing reads were first quality trimmed then assembled *de novo* using Trinity RNA-Seq (32). The assembled contigs were annotated based on similarity searches against the NCBI nucleotide (nt) and non-redundant protein (nr) databases using BLASTn and Diamond (BLASTX) (33), and an e-value threshold of 1×10^-5^ was used as a cut-off to identify positive matches. We removed non-viral hits including host contigs with similarity to viral sequences (e.g. endogenous viral elements). To reduce the risk of incorrect assignment of viruses to a given library due to index-hoping, those viruses with a read count less than 0.1% of the highest count for that virus among the other libraries were assumed to be contaminated.

### Virus Abundance and Diversity

Viral abundance was estimated using Trinity with the “align and estimate abundance” tool. The method of abundance estimation used was RNA-Seq by Expectation-Maximisation (RSEM) (34) and the alignment method used was Bowtie2 with “prep_reference” flag enabled (35). Estimated viral abundances were first standardised against the number of raw reads in each library and then compared to standardised abundances of ribosomal protein s13 (RPS13), which is stably expressed in avian hosts (21). Putative eukaryotic-associated viral transcripts were confirmed by translating open reading frames (ORFs) on Geneious Prime (v. 2022.2.1) and checked via BLASTp (https://www.geneious.com) (36). Viruses were categorised based on viral family and considered novel viruses if they shared <90% amino acid similarity within conserved regions (polymerase, viral protein (VP) regions) to known viruses.

### Virus Phylogenetic Characterisation

To infer the evolutionary relationships of virus transcripts identified, viral contigs were first translated and then combined with representative protein sequences from the same viral genus along with *Torque teno virus 1*, an outgroup from the *Anelloviridae*, obtained from NCBI GenBank. All sequences were first aligned using MAFTT (v7.4) (37) employing the E-INS-i algorithm and the alignment was trimmed using GBlocks (38, 39) to remove ambiguously aligned regions. Maximum likelihood phylogenetic trees were estimated in IQ-TREE (40) using the best fit amino acid model, LG, as determined by ModelFinder (41), with 1,000 bootstraps replicated. Phylogenetic trees were then annotated in FigTree (http://tree.bio.ed.ac.uk/software/figtree/) and Adobe Illustrator.

### Total Infectome Characterisation

Each sequencing library was evaluated for the presence and abundance of potential pathogenic bacterial, protozoal and fungal organisms known to potentially cause disease in avian species. Organism abundances were estimated as described above and summed for each bacterial, protozoal and fungal family/genus identified. Abundances were standardised against the number of raw reads within each sequencing library.

### PCR Confirmation of Viral Presence or Absence

Primers were designed to detect and confirm amplicons of yellow-eyed penguin gyrovirus using conventional polymerase chain reaction (PCR). Primers were designed using Geneious Prime^®^ 2022.2.1 (Biomatters Ltd.) from the complete genome of yellow-eyed penguin gyrovirus. In the Geneious^®^ Primer Design function, four forward primers and four reverse primers were designed with each primer 18-20 bp in length with a GC content of 42.9 - 56.0% and a T_m_ of 55.1 – 57.9°C. Primers were designed and selected based on identifying a product size of 620–820 bp with overlap and no repetition (Supplementary Table 2). PCR amplification was performed as follows: 15 minutes at 95°C; 35 cycles of 94°C for 30 seconds, 57°C for 30 seconds and 72°C for 45 seconds; 72°C for 10 minutes. Optimisation and confirmation included utilising samples with high abundances of yellow-eyed penguin gyrovirus obtained from RNA sequencing and by altering the annealing temperature by 1°C to maximise specificity.

PCR products were separated by gel electrophoresis with fluorophore (SYBR Safe DNA Gel Stain, Invitrogen, ThermoFisher Scientific) in a 1% agarose gel (Invitrogen, Life Technologies) on 80 mV for 35-45 minutes, depending on expected size and indicated quality of the PCR products, and included negative controls, positive controls and 1 kb ladder (Invitrogen, ThermoFisher Scientific) on 0.5X TBE running buffer. PCR bands were visualised by UV light using GelDoc Go Imaging System (Bio-Rad Laboratories). Bands of the expected size were excised from the successful primers, and DNA was purified using a gel cleanup kit (Qiagen QIAquick PCR & Gel Cleanup Kit). DNA was sent for bi-directional Sanger sequencing to the University of Otago Genetic Analysis Services, New Zealand. Forward and reverse sequences were pair-end assembled using

Geneious Prime^®^2022.2.1 software (version 11.0.14). Contigs were matched to known sequences on the NCBI database using the nt database using Basic Local Alignment Search Tool for nucleotides (BLASTn) (36). Contigs were confirmed with the highest hit as gyrovirus. The remainder of the samples were tested individually by tissue (lung, liver, spleen, kidney and bursa) by converting RNA to cDNA using a High Capacity cDNA Reverse Transcriptase kit (Applied Biosystems, ThermoFisher Scientific) followed by PCR using YEP GV P2-F and YEP GV P2-R primers (Table 1).

### Virus Nomenclature

Following consultation with local iwi (indigenous peoples of Aotearoa New Zealand), we propose the virus name ‘yellow-eyed penguin gyrovirus’.

## Acknowledgements

We would like to thank the New Zealand Department of Conservation staff, the Yellow-eyed Penguin Trust, Penguin Place, Penguin Rescue, and the Wildlife Hospital Dunedin. Thanks to Hamish Thompson for the illustration of the yellow-eyed penguin used in this manuscript. Thank you to local Kāi Tahu members for their collaboration and support.

## Data Availability

The novel yellow-eyed gyrovirus genomes can be found under GenBank accession numbers [pending]. Raw sequencing reads can be found in the SRA under accession numbers [pending].

## Author Contributions

Conceptualisation: J.R.W, K.J.M., K.M., J.L.G.; Data curation: J.R.W., S.H., L.B., J.D., M.B., F.J., J.L.G.; Formal analysis: J.R.W, J.L.G.; Funding Acquisition: J.R.W, K.J.M., J.L.G.; Resources: S. H., H.S.T., L.S.A., M.B., T.W., K.M., J.L.G.; Supervision: K.J.M., J.L.G.; Writing, original draft and preparation: J.R.W, J.L.G.; Writing, review and editing: J.R.W., K.J.M., S.H., H.S.T., L.S.A., T.W., J.D., F.J., M.B., L.B., E.C.H., K.M., J.L.G.

## Supporting Information Captions

**Supplementary Table 1**. Nest locations, disease status and standardised abundance of gyrovirus and PCR results from yellow-eyed penguin (*Megadyptes antipodes*) chicks (n=43) from the 2021 breeding season across sequencing libraries.

**Supplementary Table 2**. Forward and reverse primers designed for PCR testing for yellow-eyed penguin gyrovirus.

**Supplementary Figure 1**. Electron microscopy of lung tissue from a deceased yellow eyed penguin (*Megadyptes antipodes*) chick from the 2021 breeding season with respiratory distress syndrome; nucleus (N), mitochondria, vacuoles (V). Scale bars 1000 nm (left) and 500 nm (right)

**Supplementary Figure 2**. Electron microscopy of spleen tissue from a deceased yellow eyed penguin (*Megadyptes antipodes*) chick from the 2021 breeding season; nucleus (N), circular nuclear densities (C), mitochondria, vesicles (Ve). Scale bars 1000 nm (left) and 500 nm (right)

**Supplementary Figure 3**. PCR gels for identification of amplicons for the yellow-eyed penguin gyrovirus.

