## Supplementary figures and images for "A novel gyrovirus associated with a fatal respiratory disease in yellow-eyed penguin (*Megadyptes antipodes*) chicks"

### Supplementary Figure 1

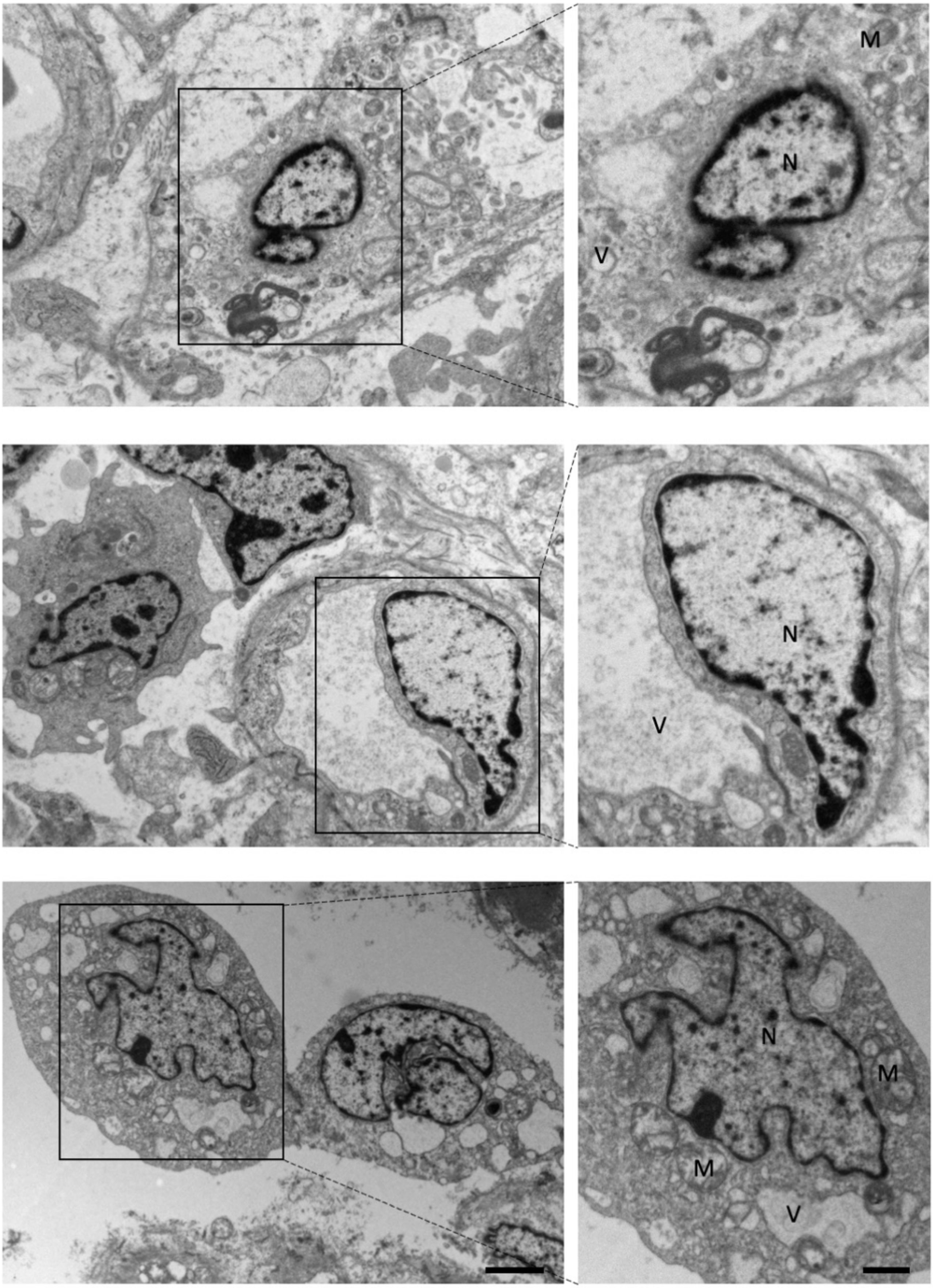

### Supplementary Figure 2

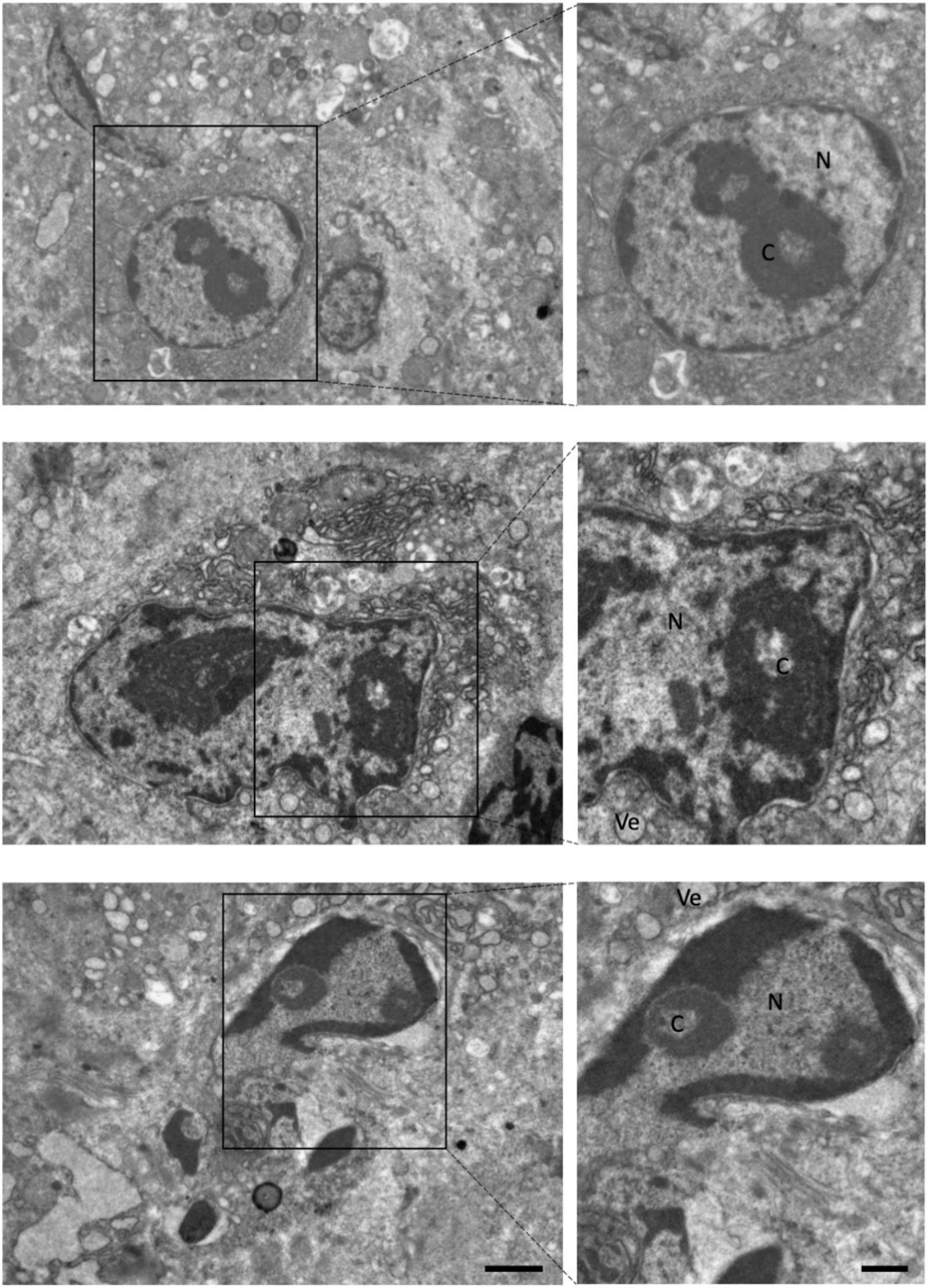

### Supplementary Figure 3

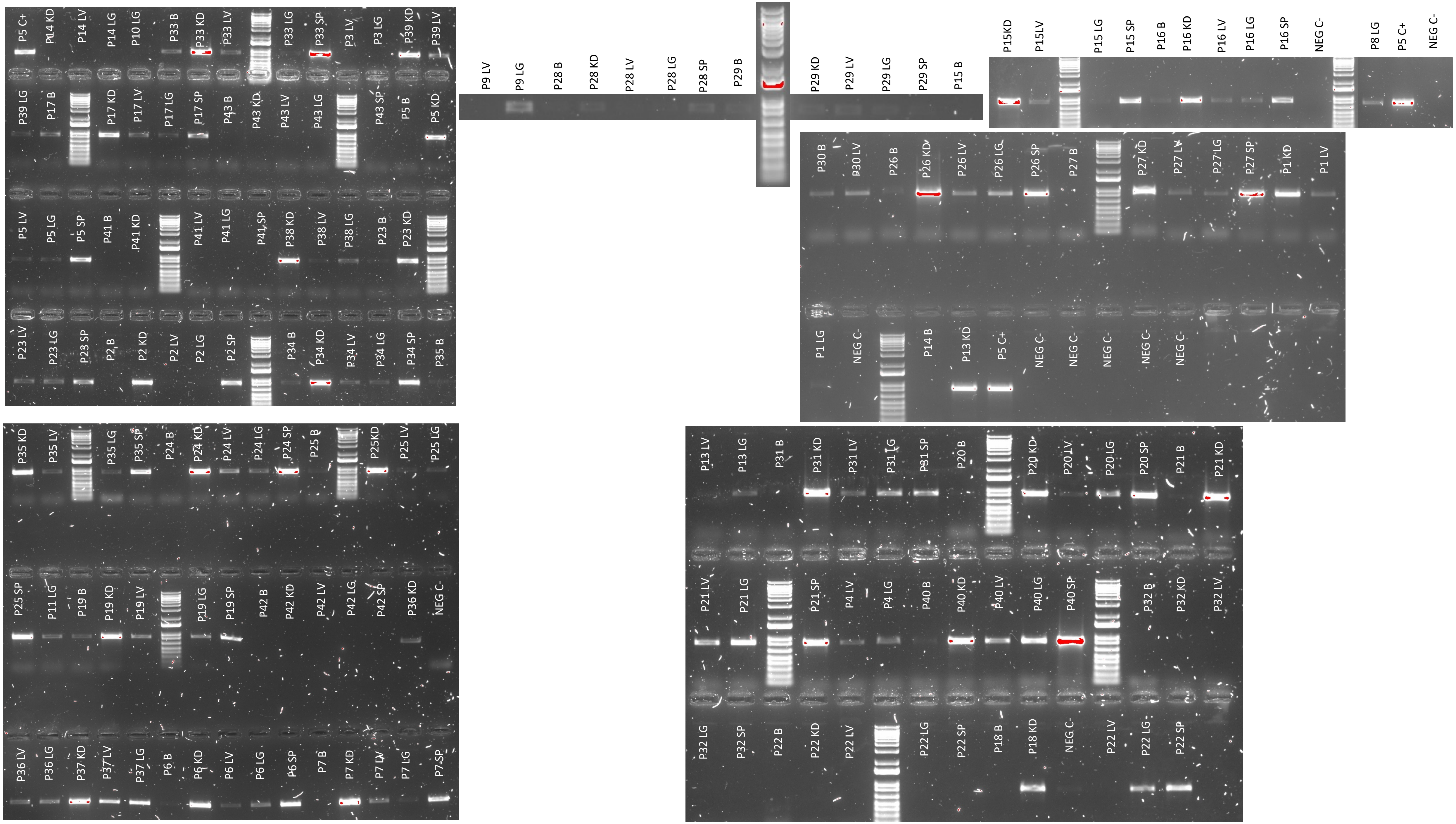
