## Supplementary Table 1 for "A novel gyrovirus associated with a fatal respiratory disease in yellow-eyed penguin (*Megadyptes antipodes*) chicks"

**Supplementary Table 1**. Nest locations, disease status, standardised abundance of yellow-eyed penguin gyrovirus and PCR results across sequencing libraries.

| **Sequencing library ID** | **No. sequencing reads** | **Sampling location** | **RDS Status** | **Standardised YEP gyrovirus abundance** | **Location of death** | **Age at death (days of age)** | **PCR results (+ positive; - negative; N/A tissue not available)** | | | | |
| --- | --- | --- | --- | --- | --- | --- | --- | --- | --- | --- | --- |
|  |  |  |  |  |  |  | **Bursa** | **Kidney** | **Liver** | **Lung** | **Spleen** |
| **P17** | 74,669,050 | North Otago | SUSPECTED | 1.716E-05 | Nest | 7 | + | + | + | + | + |
| **P3** | 58,331,101 | North Otago | CONFIRMED | 0 | Nest | 1 | N/A | N/A | - | - | N/A |
| **P33** | 63,573,823 | North Otago | NO | 1.537E-05 | Nest | 11 | + | + | + | + | + |
| **P36** | 67,376,712 | North Otago | NO | 6.555E-05 | Nest | 10 | N/A | + | + | + | N/A |
| **P37** | 55,923,975 | North Otago | SUSPECTED | 1.869E-05 | Nest | 9 | N/A | + | + | + | N/A |
| **P38** | 58,688,665 | North Otago | CONFIRMED | 0.0001405 | Nest | 3 | N/A | + | + | + | N/A |
| **P39** | 81,514,743 | North Otago | CONFIRMED | 3.387E-05 | Nest | 14 |  | + | + | + | N/A |
| **P41** | 85,110,255 | North Otago | NO | 0 | Nest | 15 | - | - | - | - | - |
| **P42** | 53,475,625 | North Otago | CONFIRMED | 0 | Transfer | 7 | - | - | - | - | - |
| **P43** | 72,248,283 | North Otago | CONFIRMED | 0 | Nest | 9 | - | - | - | - | - |
| **P5** | 73,258,778 | North Otago | CONFIRMED | 2.197E-05 | Nest | 10 | - | + | + | + | + |
| **P6** | 53,782,568 | North Otago | CONFIRMED | 7.238E-06 | Hospital | 5 | + | + | + | + | + |
| **P7** | 57,602,812 | North Otago | CONFIRMED | 6.363E-06 | Hospital | 6 | - | + | + | + | + |
| **P1** | 62,189,455 | Otago Peninsula | CONFIRMED | 5.335E-06 | Hospital | 4 |  | + | + | + | N/A |
| **P10** | 64,312,113 | Otago Peninsula | CONFIRMED | 6.22E-08 | Hospital | 10 | N/A | N/A | N/A | - | N/A |
| **P12** | 61,939,914 | Otago Peninsula | CONFIRMED | 5.651E-07 | Hospital | 2 | - | N/A | N/A | N/A | N/A |
| **P13** | 70,571,497 | Otago Peninsula | SUSPECTED | 8.607E-06 | Hospital | 3 | N/A | + | + | + | N/A |
| **P14** | 62,826,954 | Otago Peninsula | CONFIRMED | 4.654E-07 | Hospital | 5 | + | - | - | - | N/A |
| **P20** | 75,276,121 | Otago Peninsula | SUSPECTED | 6.707E-06 | Hospital | 4 | - | + | + | + | + |
| **P21** | 62,688,595 | Otago Peninsula | CONFIRMED | 2.342E-05 | Hospital | 3 | - | + | + | + | + |
| **P22** | 70,999,561 | Otago Peninsula | SUSPECTED | 0 | Euthanised in hospital | 3 | - | - | - | - | - |
| **P24** | 61,977,039 | Otago Peninsula | SUSPECTED | 1.873E-05 | Hospital | 5 | - | + | + | + | + |
| **P25** | 61,458,622 | Otago Peninsula | CONFIRMED | 9.305E-05 | Hospital | 5 | + | + | + | + | + |
| **P31** | 44,577,939 | Otago Peninsula | CONFIRMED | 1.872E-05 | Hospital | 3 | - | + | + | + | + |
| **P32** | 62,255,608 | Otago Peninsula | CONFIRMED | 0 | Euthanised in hospital | 3 | - | - | - | - | - |
| **P34** | 61,307,927 | Otago Peninsula | CONFIRMED | 5.308E-05 | Euthanised in hospital | 8 | + | + | + | + | + |
| **P35** | 65,619,249 | Otago Peninsula | CONFIRMED | 1.491E-05 | Euthanised in hospital | 7 | + | + | + | + | + |
| **P4** | 58,506,567 | Otago Peninsula | CONFIRMED | 1.333E-06 | Hospital | 8 | N/A | N/A | + | + | N/A |
| **P40** | 53,128,986 | Otago Peninsula | CONFIRMED | 2.116E-05 | Hospital | 4 | + | + | + | + | + |
| **P11** | 67,686,096 | The Catlins | CONFIRMED | 1.477E-08 | Hospital | 6 | N/A | N/A | N/A | + | N/A |
| **P15** | 66,316,575 | The Catlins | CONFIRMED | 2.625E-05 | Hospital | 6 | + | + | + | + | + |
| **P16** | 54,475,176 | The Catlins | CONFIRMED | 6.947E-05 | Hospital | 6 | + | + | + | + | + |
| **P18** | 56,020,312 | The Catlins | CONFIRMED | 5.052E-06 | Hospital | 9 | - | + | + | + | + |
| **P19** | 79,229,715 | The Catlins | CONFIRMED | 1.215E-05 | Hospital | 5 | + | + | + | + | + |
| **P2** | 44,986,520 | The Catlins | CONFIRMED | 1.672E-05 | Hospital | 5 | - | + | - | - | + |
| **P23** | 62,703,338 | The Catlins | CONFIRMED | 1.97E-05 | Euthanised in hospital | 6 | + | + | + | + | + |
| **P26** | 74,439,794 | The Catlins | CONFIRMED | 1.635E-05 | Hospital | 5 | + | + | + | + | + |
| **P27** | 73,492,561 | The Catlins | CONFIRMED | 9.688E-06 | Hospital | 5 | - | + | + | + | + |
| **P28** | 65,849,276 | The Catlins | CONFIRMED | 0 | Hospital | 9 | + | - | - | + | + |
| **P29** | 54,587,920 | The Catlins | CONFIRMED | 0 | Hospital | 6 | + | + | + | - | - |
| **P30** | 54,285,374 | The Catlins | SUSPECTED | 1.23E-05 | Hospital | 5 | + | N/A | + | N/A | N/A |
| **P8** | 46,733,551 | The Catlins | SUSPECTED | 0 | Nest | 4 | N/A | N/A | N/A | + | N/A |
| **P9** | 64,500,121 | The Catlins | SUSPECTED | 3.256E-07 | Nest | 4 | N/A | N/A | + | + | N/A |
