## Supplementary Table 2 for "A novel gyrovirus associated with a fatal respiratory disease in yellow-eyed penguin (*Megadyptes antipodes*) chicks"

**Supplementary Table 2**.  Forward and reverse primers designed for PCR testing.

| **Primers** | **Nucleotide sequence (5’  3’)** | **Size (base pairs)** |
| --- | --- | --- |
| YEP GV P1-F | CCCCTTTCAAGCAGAAGAC | 749 |
| YEP GV P1-R | GAAGAAGAAGACGGTGGCA |  |
| YEP GV P2-F | CTGGATCCGATACAGACCTTG | 842 |
| YEP GV P2-R | CATCTCTTACCGGATGATAGAG |  |
| YEP GV P3-F | CTATTTTGGCTGGGGAACC | 704 |
| YEP GV P3-R | CCTACCGTAATAAGTGGGCTT |  |
| YEP GV P4-F | AAAGTGTTCAGCAAATGGCAG | 640 |
| YEP GV P4-R | GCATGTGTGTAGATCTCGGT |  |
